# Retrosplenial cortical representations of space and future goal locations develop with learning

**DOI:** 10.1101/315200

**Authors:** Adam M. P. Miller, William Mau, David M. Smith

## Abstract

The retrosplenial cortex (RSC) is important for long-term contextual memory and spatial navigation, but little is known about how RSC neural representations develop with experience. We recorded neuronal activity in the RSC of rats as they learned a continuous spatial alternation task and found that the RSC slowly developed a population-level representation of the rat’s spatial location and current trajectory to the goal. After the rats reached peak performance, RSC firing patterns became predictive of navigation accuracy and even began to represent the upcoming goal location as the rats approached the choice point. These neural representations emerged at the same time that lesions impaired performance, suggesting that the RSC gradually acquired task representations that contribute to navigational decision-making.

---

Long-term memory is thought to rely on the neocortex and is critical for a variety of cognitive processes, including attention, decision making, and new learning (1-3). In spatial navigation, for example, subjects use long-term memory for the spatial layout of the environment—including routes, goal locations, and task rules—to achieve their navigation goals (4). Much of what we know about spatial memory and navigation comes from work on the hippocampus (5). However, many spatial memories do not require the hippocampus after they have become well-learned (6-8). In contrast, the retrosplenial cortex (RSC) may play an important role in long-term spatial memory (9).

The RSC is closely interconnected with brain regions known to play a role in navigation, including the hippocampus and anterior thalamus (10, 11). Recent findings suggest that long-term contextual memories depend on the RSC (12-14), and that RSC damage impairs navigation in humans and rodents (15-18). An extensive fMRI literature similarly suggests an RSC role in spatial memory (19-21), including in the representation of spatial heading (22, 23) and landmarks (24, 25). Furthermore, individual RSC neurons exhibit spatially localized firing (26, 27) and directional firing (26, 28, 29), and they respond strongly to navigational cues (30). Recent work has also shown that RSC neuronal activity is modulated by allocentric, egocentric, and route-centered spatial reference frames (31, 32), consistent with the idea that the RSC is well positioned to integrate information from each of these domains (33).

Despite this growing literature, surprisingly little is known about how RSC neuronal ensembles might represent spatial information. For example, despite suggestive evidence from several studies (26, 27, 31, 32), it is not clear whether the RSC generates a map-like representation that indicates the subject’s position within the environment, whether RSC representations support spatial memory functions, or whether spatial representations emerge immediately or develop as a function of repeated experience. To examine this, we trained rats on a memory guided spatial navigation task and we examined RSC ensemble representations across stages of learning.

## Results

We recorded from 637 RSC neurons in 12 rats as they learned and performed a continuous spatial alternation task on a modified T-maze (**Fig. 1a**). Rats were given daily training sessions until they reached a criterion of 90% correct, followed by at least four additional asymptotic performance sessions. Our recordings targeted the granular b subregion of the RSC bilaterally, although small numbers of neurons from the granular a subregion and the dysgranular RSC were also included (**Supplementary Fig. 1**). There were no conspicuous differences in the firing properties of neurons recorded in different subregions, different hemispheres, or at different AP coordinates. We restricted our analyses to training days that were common to all subjects. This typically included the first, middle, criterial, and post-criterial sessions (i.e. overtraining), but in some cases included additional days (see **Supplementary Fig. 2** for details).

**Figure 1.**
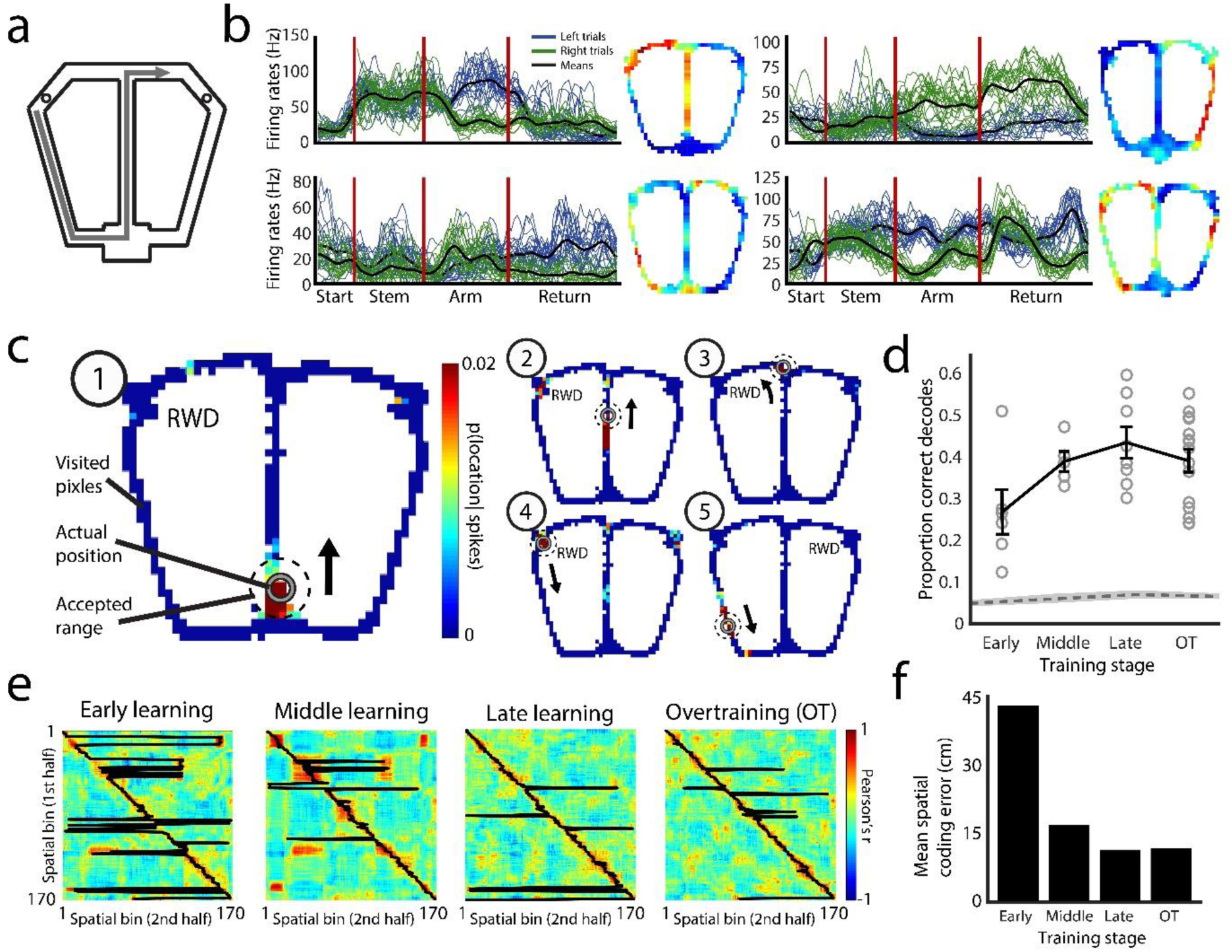
Neural populations in the retrosplenial cortex (RSC) developed a representation of the maze with learning. (**a**) Schematic diagram of the continuous T-maze alternation task. After visiting one of the two reward locations (circles), rats returned to the stem and had to approach the opposite location for reward. (**b**) RSC neuronal firing reliably distinguished between maze locations from one trial to the next. The firing of four example neurons illustrating the trial by trial firing as rats traversed the maze on left and right trials (see also (31)). Red vertical lines show maze section boundaries. Mean firing rate maps show the spatial distribution of firing rates for each example neuron. Note that neurons typically fired over large areas of the maze and despite considerable variability, differences in firing rates between maze regions were quite reliable. (**c**) A Bayesian decoder was applied to the activity of simultaneously recorded ensembles of RSC neurons to predict the rat’s current location on the maze based on instantaneous firing. Five decoded instances from one trial are shown. Colored pixels in the shape of the maze indicate the probability of the rat being in that pixel given the instantaneous spiking activity, with warmer colors corresponding to higher probabilities. The highest probability pixel was taken as the decoded location. The rat’s actual location is shown by the grey circle, and the rat’s current direction of travel is indicated by the black arrow. Decoded locations falling within the dashed circle were counted as correct. (**d**) RSC ensembles show improved encoding of spatial position with training. The decoder’s success rates for individual sessions are plotted as open circles, while the mean for each training stage is shown by the line plot +/- SEM. Decoding improved significantly with training and was always far more accurate than was expected by chance (gray area shows the center 95% of the shuffle distribution). (**e**) Improvements in spatial representation were not due to learning-related changes in behavior. A strict behavioral criterion was applied to remove instances of atypical behavior (see **Supplementary Fig. 3**). A full lap around the maze was divided into 170 spatial bins (3 cm per bin) and correlations were computed between firing rate vectors from the first and second half of each session at all spatial bins. Correlation matrices from early, middle, and late learning sessions, and from overtraining, show correlations between firing occurring in each bin during the first half and second half of sessions. The black line connects the pixels of highest correlation between the two session halves at each spatial bin. Deviations from the diagonal are indicative of spatial coding errors. (**f**) Mean spatial coding error over all bins is plotted for each learning stage.

### RSC neural populations develop a representation of the maze with learning

Many neurons in the RSC showed reliable firing rate differences over the spatial extent of the maze (**Fig. 1b**). Consistent with previous reports (26, 27, 31, 32, 39), these neurons exhibited larger firing fields with higher background firing rates than those of hippocampal place cells (27, 40). Nonetheless, we found that RSC ensembles developed a reliable representation of the maze as rats learned the alternation task. To quantify this, we used Bayesian decoding to determine whether we could predict the rat’s current spatial location solely on the basis of the activity of the recorded neural ensemble (41, 42) (**Fig. 1c**). We found that the rat’s position could be accurately predicted at a rate far greater than chance even during earliest stages of learning (p < 0.001, compared to a control distribution generated by shuffling firing rates across time bins, see Methods; **Fig. 1d**), and that decoding accuracy improved significantly as the rats learned (F(3, 30) = 3.02, p < 0.05). At asymptote, the rat’s position could be accurately predicted within 4.5 cm about 40% of the time.

To confirm that this improvement in representation was not attributable to learning-related changes in behavior (e.g., more stereotyped maze running as the rats became more practiced), we employed a correlational reconstruction technique and we limited the analysis to trials in which the rat followed a stereotyped path (see Methods and **Supplementary Fig. 3**). We combined neurons from all rats into a single population for each training stage and calculated mean firing rate vectors for each spatial bin (170 spatial bins forming a full lap around the maze; each bin included 3cm of track) separately for the first and second halves of each session. Mean ensemble firing rate vectors from the first and second halves of the session were correlated and plotted as correlation matrices (**Fig. 1e**). If spatially localized firing patterns were reliable across session halves, the highest correlation should occur between visits to the same location (along the diagonal), while deviations from the diagonal indicate instances of unreliable spatial coding (i.e. spatial coding errors). Consistent with the Bayesian analysis above, spatial coding was far more reliable and accurate than expected by chance at all stages of learning (all p < 0.001, compared to a distribution generated by shuffling first-half/second-half neuron pairings) and the representation improved with learning, as indicated by a 70% reduction in spatial coding errors from early learning to overtraining (p < 0.005, compared to a distribution generated by shuffling neurons between stages; **Fig. 1f**). Together, these two independent analysis methods, Bayesian decoding and correlational reconstruction, provide compelling evidence that representations of the maze emerged in the RSC with learning.

### RSC neural populations develop trial-type specific responses on the stem of the maze

In the continuous alternation task, rats must use memory for the previous (or upcoming) reward location to know where to go on the current trial. This memory might be encoded, in part, by differential firing as rats traverse the stem of the maze on the approach to the choice point (43). We found that RSC neurons exhibited two distinct firing patterns on the stem of the maze depending on whether the rat would later turn left or right (i.e. trial-type specific firing; **Fig. 2a**). At asymptote, 21% of the neurons showed trial-type specific firing that was significantly greater than expected by chance (p < 0.001, compared to distributions generated by shuffling left and right trial types; **Fig. 2b**, **Supplementary Fig. 5a**). It is unlikely that these firing patterns were caused by small differences in the rat’s behavior as they traversed the stem, as neither head direction (r = −0.10, p = 0.10), nor lateral position (r= −0.04, p = 0.46), nor running speed (r = 0.06, p = 0.35) was significantly correlated with trial-type specific firing (**Supplementary Fig. 5b-d**).

**Figure 2.**
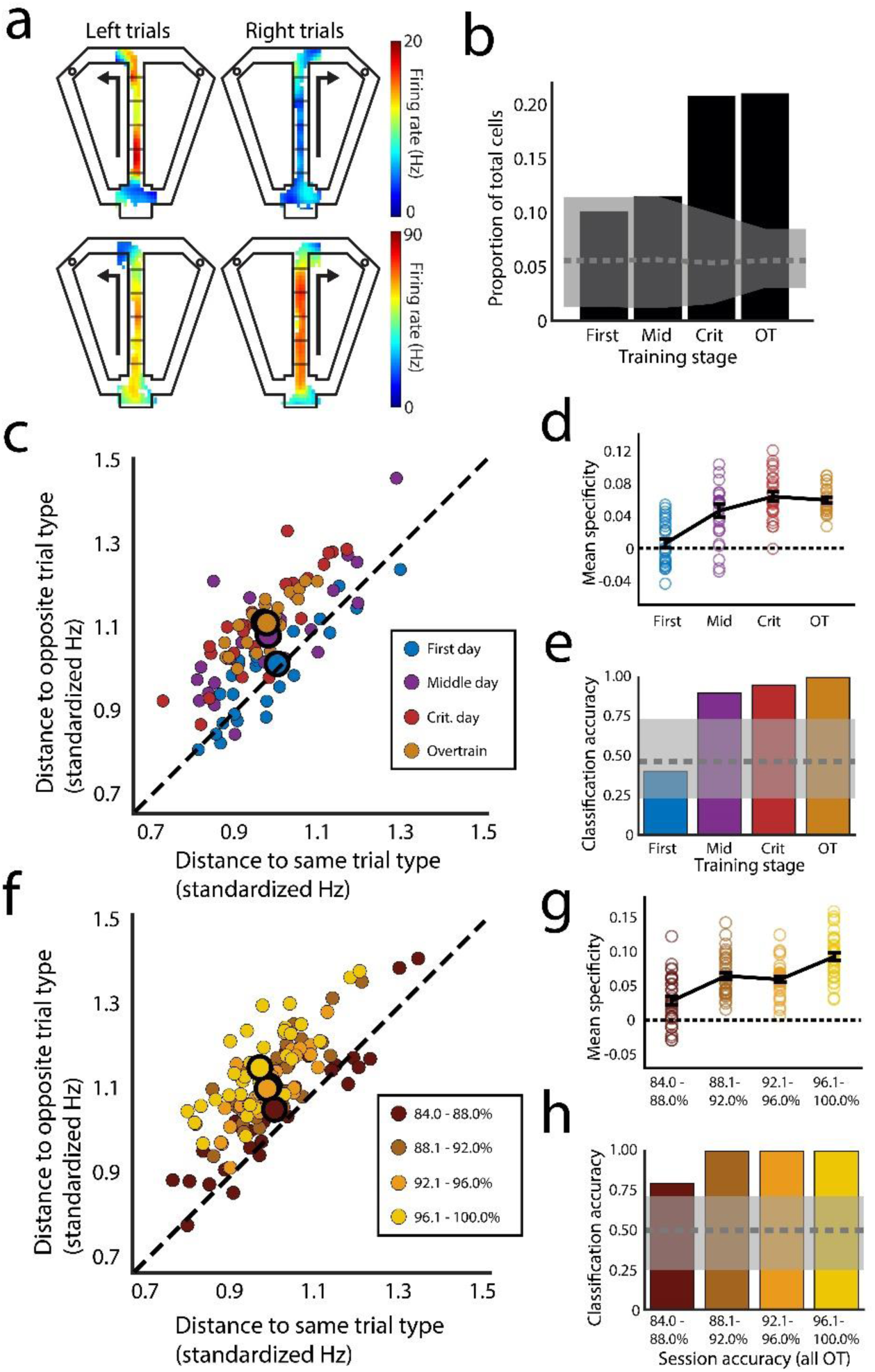
Neurons in the RSC develop trial-type specific firing on the stem. (**a**) Two examples of RSC neurons (rows) that fired differently on the stem depending on whether the rat was about to turn left or right (i.e. trial-type specific firing). Firing rate maps are shown for left and right turn trials, with the analyzed sectors of the stem indicated. (**b**) The proportion of RSC neurons showing trial-type specific firing is plotted for the first (First), middle (Mid), and last (criterion, Crit) learning day, and for all overtraining (OT) days. Gray shading shows the chance level on each day. (**c**) Trial-type specificity of RSC neural ensemble firing increased with training. Each colored dot shows ensemble activity from one trial (combined across subjects and sessions, see Methods) plotted in terms of its distance from the mean of the same and opposite trial types. Points along the dotted line are equidistant to both trial types, indicating that the ensemble showed no preference for left or right trials, while points farther from the dotted line indicate stronger ensemble preferences for one trial type over the other. Large dots outlined in black illustrate the mean for each learning stage. Note that ensemble activity diverges from the unity line as learning progresses. (**d**) Trial-type specificity of the RSC ensemble increased with training and was greater than chance by the middle training day. Individual trials (small dots from **c**) are plotted as open circles, with the mean for each training stage illustrated by the line plot +/- SEM. (**e**) The ability to classify trials (left or right) solely on the basis of ensemble firing patterns improved with learning, from chance (gray shading) on the first day of training, to perfect accuracy during overtraining. (**f-h**) The trial-type specificity of RSC ensemble firing was greater during sessions with better alternation performance. Plots are the same as **c-e,** except that all data were taken from overtraining sessions that were grouped according to behavioral performance (% correct choices for the session). Note that ensemble activity shows increased trial-type specificity and improved classification of left and right trials during sessions with superior behavioral performance.

To determine how these responses were related to behavior, we combined neurons from all rats into a single ensemble for each training stage and then calculated the similarity between ensemble responses occurring on the stem during left and right trials. We found that RSC activity became more trial-type specific with learning (F(3, 100) = 20.02, p < 0.001, **Fig. 2c,d**), indicating that activity on left trials became more similar to other left trials, and less similar to right trials (and likewise with right trials). Ensemble activity also became more predictive of turning behavior with training. We classified trials as either left or right depending on whether ensemble activity occurring on the stem was more similar to the mean activity of left or right trials, and found that classification accuracy increased with learning (p < 0.05, compared to a control distribution generated by shuffling neurons between training stages, **Fig. 2e**), from chance classification on the first day (46.15%) to perfect classification during overtraining.

We also found that ensemble activity became more predictive of accurate navigation. We binned the overtraining sessions into four categories based on performance, and found that sessions with better behavioral performance had higher neural trial-type specificity (F(3, 132) = 24.38, p < 0.001; **Fig. 2f,g**). Trial-type specificity was three-fold higher in the sessions with the best performance (96.1-100% accuracy) than in sessions with poor performance (84%-88% accuracy; t(66) = 7.68, p < 0.001). Similarly, our ability to classify trials as left or right was better on superior performance days than on poor performance days (p < 0.05, compared to a distribution generated by shuffling neurons between performance groupings; **Fig. 2h**). This is the first direct evidence that trial-type specificity is associated with good performance in a spatial memory task.

### RSC ensembles represent future goal locations

In human subjects, the RSC is active during navigational route planning (44), and, in rodents, a similar process is associated with forward-sweeping simulations of possible trajectories to the goal in the hippocampus (45). We were therefore interested in determining whether the RSC similarly represents distant goal locations before the rats turned left or right. To do this, first confirmed that the RSC distinctly represented the two reward locations by combining neurons across subjects and comparing ensemble activity at the reward locations on left and right trials. We found that RSC activity was more similar during visits to the same reward location than between visits to opposite locations, and that these patterns became more distinct as the rats learned (F(3,100) = 4.34, p < 0.01, **Supplementary Fig. 4**, see also (30)).

To then test whether RSC populations represented the distant goal locations while the rat was on the stem, we used Bayesian decoding to compute probability distributions reflecting the predicted location of the rat given ensemble firing activity occurring on the stem (**Fig. 3a-c;** see Methods for details). We found that a sizeable portion (25.86%) of the decoded probability distribution was located in the reward areas far ahead of the rat’s actual location (reward areas are shown in **Fig. 3b**). Decoding to the reward areas increased with learning (F(3, 30) = 2.95, p = 0.05, **Fig. 3c,d**) from marginally greater than chance during the early and middle stages of learning (early, t_(5)_=2.22, p = 0.08; middle, t_(4)_ ^=^ 2.53, p = 0.06) to far greater later in learning (late, t_(7)_ = 4.76, p < 0.005; overtraining, t_(14)_ = 9.21, p < 0.001). These representations included both punctate instances of unambiguous decoding to the reward area and times when the firing patterns momentarily became more similar to the firing patterns observed when the rat was at the reward. Similar results were obtained when the analysis was limited to the proportion of absolute classifications rather than the proportion of the decoded probability distribution.

**Figure 3.**
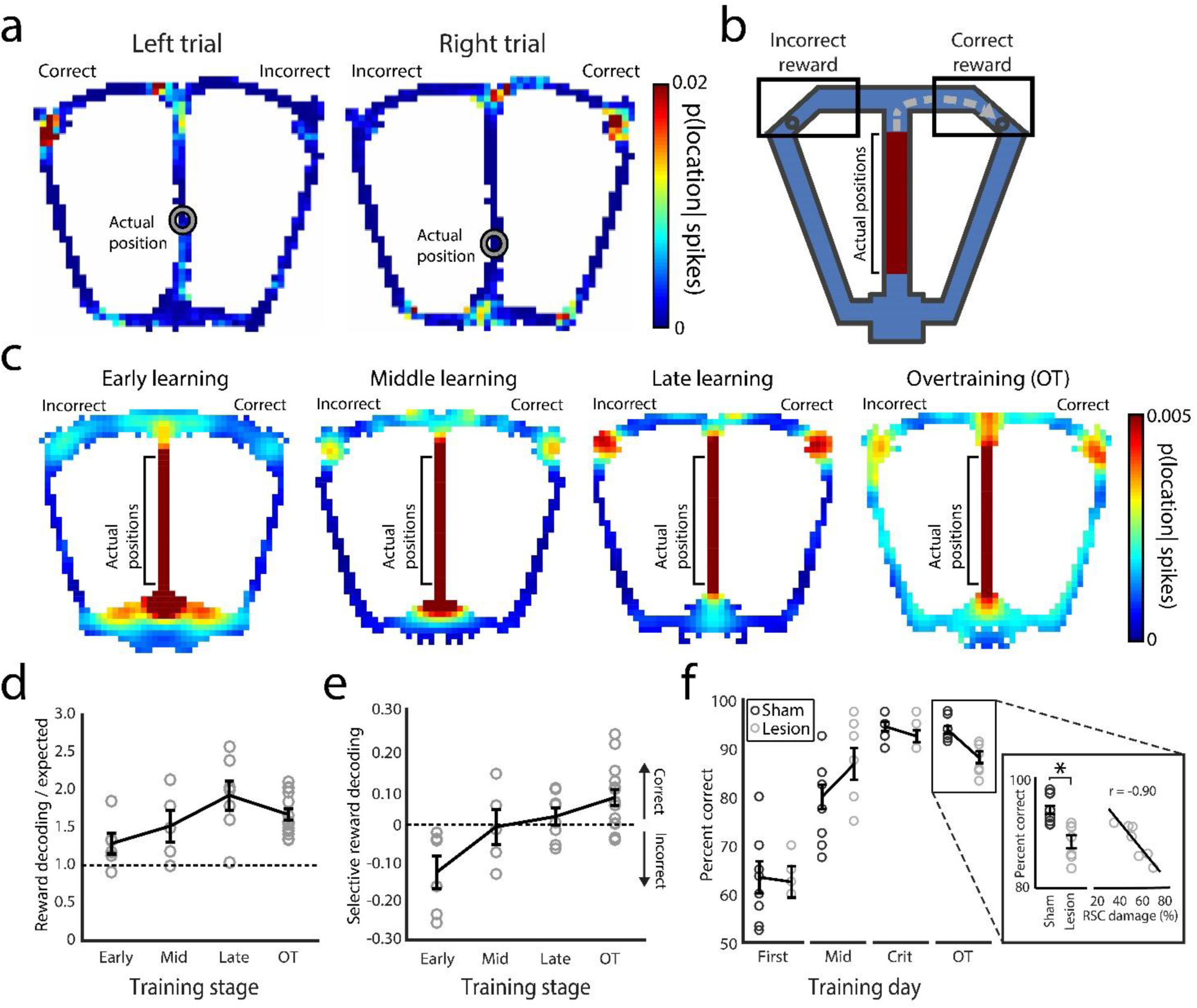
RSC ensembles represent upcoming reward locations. (**a**) Bayesian decoding was used to identify representation of the reward locations as rats approached the choice point. Two examples are shown, from left and right trials, of decoded instances when ensemble firing patterns were more consistent with the upcoming reward area than the rat’s actual position (gray circle). (**b**) The analyses of decoded spatial information focused on the two reward locations and the distal part of the goal arms approaching each reward (black rectangles) but, importantly, was limited to time windows when the rat was located on the stem (red). (**c**) Heat maps illustrating the average decoded probability from all of the 200 ms time bins used for decoding as the rat traversed the stem, with separate heat maps shown for each learning stage. For illustration purposes, the data from the left trials are mirror reversed so that all the data are shown with the correct goal location to the right and the incorrect (previous) goal location shown to the left. Note the faint clouds of probability at the reward areas (i.e. decoding to the reward areas) during the early learning stage. This becomes more prominent through late learning and only becomes selective for the correct reward area during overtraining. Stem locations are uniformly red because the decoding is most prevalent at the rat’s actual current location on the stem. (**d**) Decoding to the two reward areas increased with training and surpassed chance levels (uniform non-local decoding, dashed line, see Methods) only at criterion and during overtraining. Reward decoding was calculated by dividing the decoded probability that the rat was located in the rectangles of **b** by the expected probability as determined by relative area. Individual ensembles are plotted as open circles, while average reward area decoding for each training stage is shown by the line plot +/- SEM. (**e**) Selective decoding to the correct reward area only emerged as statistically significant during overtraining. Future reward decoding was defined as the normalized difference between decoding to the correct reward area and the opposite (incorrect) reward area ((p(correct) – p(incorrect)) / p(correct) + p(incorrect)). Individual ensembles are plotted as open circles, while average reward area decoding for each training stage is shown by the line plot. (**f**) Permanent lesions of the retrosplenial cortex selectively impaired spatial alternation performance after learning. Behavioral performance is plotted for the first (First), middle (Mid), and last (criterial, Crit) learning days, and asymptotic performance days (overtraining, OT). Performance for each control and lesion rat (left) is shown as open circles, with the mean indicated by line plots +/- SEM. The inset illustrates overtraining performance, along with the correlation between performance and lesion size.

RSC ensembles initially represented both the correct and incorrect reward areas equally. However, they began to preferentially represent the correct reward area when the rats became proficient at the task (i.e. during overtraining). To quantify this, we compared the decoded probability that the rat was located in the correct reward area, p(correct), with the opposite reward area, p(incorrect, **Fig. 3b**), and found that RSC ensembles began to preferentially represent the correct reward area with learning (F(3, 30) = 7.68, p < 0.001, **Fig. 3c,e**). Early in learning, RSC populations showed a numerical preference for the incorrect (previously visited) reward area. However, after correcting for multiple comparisons, we only found a significant preference between the two reward areas during overtraining sessions (t_(14)_ = 3.36, p < 0.005). This preference for the future reward area during overtraining was not correlated with differences in head direction (r = 0.05, p = 0.85), lateral position (r = 0.11, p = 0.69) between left and right trials, or running speed (r = 0.22, p = 0.43; **Fig. 3f**).

The slow development of task responses in the RSC with learning is consistent with previous studies showing that the RSC plays a selective role late in learning (19, 46, 47). We assessed the involvement of the RSC in the continuous alternation task using neurotoxic lesions of the RSC in a separate group of rats and found that performance was impaired during the same learning stage that representations of the correct goal location emerged in our neural data. A two-way repeated measures ANOVA comparing the performance of control rats and rats with neurotoxic (NMDA) lesions of the RSC over learning revealed a significant training stage by lesion group interaction (F_(6, 90)_ = 2.832, p < 0.05, **Fig. 3f; Supplementary Fig. 6a**). Post-hoc comparisons confirmed that the impairment was selective to overtraining ^performance^ (t_(15)_ = 3.47, p < 0.005; **Fig. 3f inset left**) and tightly correlated with lesion size (r = −0.90, p < 0.01; **Fig. 3f inset right**).

## Discussion

These findings provide the first direct evidence that rich and detailed spatial representations develop in the RSC over the course of learning, including information about the rat’s current spatial location, the current trajectory, reward locations, and even simulations of upcoming goal locations. In contrast to the rapid spatial coding seen in the hippocampus, many RSC representations developed slowly as the rats learned the task and some representations, such as simulations of future goal locations, did not emerge until after learning. These results suggest that the RSC is a key component of the neocortical system for storing long-term spatial memory, consistent with recent molecular and optogenetic work (13, 14, 18).

In addition to encoding the rat’s current spatial position, we also found that RSC firing patterns predicted the rat’s future navigation behavior, as indicated by firing patterns that were specific to the left and right trials as the rats traversed the stem. This is consistent with numerous reports of trajectory specific firing in the hippocampus (43, 48, 49). In the RSC, previous studies have shown that RSC firing is sensitive to current trajectory (30) and the sequences of turns that define that trajectory (31, 32), although these tasks did not involve a memory demand. Here, we found that trial-type specific firing is also prevalent in a memory-guided navigation task and we show for the first time that population-level representations were related to task performance, with greater trial-type specificity associated with greater choice accuracy. Together, these findings suggest that accurate alternation performance may be supported, in part, by distinct RSC representations of the two trajectories, left-to-right and right-to-left, that the subject must take to arrive at the correct reward location for each trial. If so, this may reflect an RSC contribution to spatial working memory (50, 51).

We observed that goal locations were a prominent component of the RSC representations in our data, consistent with our previous work (27, 30). In addition to showing that RSC activity occurring at each of the goal locations became more distinct with training, we also found that RSC ensembles transiently represented the goal locations as the rats traversed the stem of the maze and approached the choice point. Similar representations have been observed in other brain regions including the hippocampus and the ventral striatum, where simulations took the form of sequential reactivation of neurons encoding locations along the path immediately ahead of the rat (45) and the reactivation of reward-encoding neurons (52, 53), respectively. Our data contain elements of each of these findings, with RSC populations reactivating spatial firing patterns corresponding to locations far ahead of the rat along the path to the reward as well as frequent instances of firing patterns that specifically corresponded to the reward location. However, the emergence of these representations in our data set followed a distinct temporal profile in that they were initially absent during learning and only become apparent after the rat was well-trained. Selective representations of the rat’s future goal did not appear until after other RSC representations (e.g., representations of the reward locations) were fully formed. The late appearance of these responses may reflect the development of a neural schema capable of supporting goal-directed behavior (3, 36).

The observation of future reward simulation suggests an RSC role in memory and planning, which is consistent with studies of the default mode network. This network, which includes the RSC, prefrontal cortex, posterior parietal cortex and hippocampus, mediates the constructive memory processes that underlie both episodic memory and the ability to imagine future events and situations (2, 38). Notably, many of these regions are also involved in route planning in humans (15, 16, 44, 54), and evidence consistent with the representation of future routes and goal locations has been reported in several of the same regions in rats (40, 45, 55, 56). As other authors have noted, these relatively simple representations seen in rodents may serve as rudimentary building blocks for the more complex kinds of future simulation seen in human subjects (57, 58). In our data, simulations of both goal locations emerged as the rats learned, and they only became selective for the correct location after the rat reached asymptotic levels of performance. Together with our finding that RSC lesions specifically impaired task performance at asymptote, these data suggest that future reward simulations in the RSC contribute to route planning, but only after other RSC spatial representations have become sufficiently stable.

The factors that drive the development of RSC spatial representations are not known. However, the functional similarities and anatomical connectivity between the hippocampus and RSC (9) suggest that the slow emergence of RSC representations may reflect consolidation of information from the hippocampus (59, 60). Consistent with this idea, contextual fear memories depend on the hippocampus early after learning but later become more reliant on the RSC (13, 61). In one particularly striking example, optogenetic reactivation of an RSC context representation was sufficient to evoke a contextual fear memory, even when the hippocampus was inactivated (14). Systems consolidation theory holds that spatial and episodic memories, which initially depend on the hippocampus, are eventually transferred to distributed cortical representations (59, 60) and several of our observations are consistent with that idea. RSC lesions had no discernable effect on the early stages of learning and only impaired performance after the task was well learned. Moreover, the performance deficit, though modest, was remarkably well correlated with the amount of tissue damage and spatial representations were spread across a wide swath (~5mm) of cortex with no obvious functional segregation, consistent with a widely distributed representation. These representations persisted for the duration of recording, up to 30 days beyond the first exposure to the maze in some subjects, which is consistent with long-term memory, although their stability has not been assessed over longer durations. The cortical memory representations that support spatial navigation likely extend beyond the RSC, to include other midline cortical regions such as the anterior cingulate (62) and prefrontal cortex (40). Indeed, these regions may support the relatively good alternation performance seen in subjects with RSC lesions, possibly reflecting compensatory processes associated with permanent lesions (63). However, the complex spatial representations seen in the RSC, which have not been observed in other cortical regions (40), suggest that examination of the RSC and its interactions with the hippocampus and other memory regions of the brain will be particularly important for understanding spatial cognition and memory more generally.

## Author Contributions

A.M.P.M. and D.M.S. designed the experiments and wrote the paper. A.M.P.M. and W.M. conducted the experiments. A.M.P.M. performed the analyses.

## Acknowledgements

We thank Sarah Parauda, Keunhyung Yu, Alexandra Tse, and Hui Jun Li for assistance with animal training and electrode hyperdrive fabrication. We thank Howard Eichenbaum and A. David Redish for helpful comments on the manuscript. This work was supported by an NIH grant R01 MH083809 to D.M.S.

## Methods

### Subjects

Subjects were 32 adult male Long Evans rats (Charles River Laboratories, Wilmington, MA) weighing 250g-300g upon arrival. Twelve rats were used in the neurophysiology study and 20 rats were used in the lesion study. Of the 10 rats that received RSC lesions, 2 were excluded from the analysis due to hippocampal damage, and 1 was excluded because the RSC damage was unilateral. Rats were placed on a 12hr/12hr light/dark cycle with lights on at 8am and allowed to acclimate to the vivarium for at least one week prior to surgery. After recovery from surgery, rats were placed on food restriction until they reached 80-85% of their free-feeding weight. Water was always available ad libitum. All procedures complied with the guidelines of the Cornell University Animal Care and Use Committee.

### Surgery

#### Neurophysiology

Fifteen rats had a custom-built electrode microdrive implanted, which contained 20 moveable tetrodes (16 recording tetrodes and 4 reference tetrodes) made from twisting four 17μm platinum/iridium (90%/10%) wires, platinum plated to an impedance of 100- 500 kΩ, and arranged in two 10-tetrode linear arrays (one in each hemisphere) that spanned approximately 5mm along the rostrocaudal axis of the brain. Tetrodes were stereotaxically positioned bilaterally just beneath the cortical surface (2-7 mm posterior to Bregma, ±1.5mm lateral) with the tetrodes angled 30 degrees toward the midline. Rats were given 7 days to recover from surgery prior to lowering the tetrodes into the RSC (35-70 μm daily) over the course of several days until a depth of at least 1 mm was reached to ensure that the tetrodes were in the granular b subregion (discussed below).

#### Lesions

Twenty rats were anethetized with isoflorane gas (1-5% in oxygen) and placed in a Kopf stereotaxic apparatus. The skin was retracted and holes were drilled through the skull above each of the injection sites. Ten rats received bilateral neurotoxic (N-methyl-D-aspartate [NMDA], 10μg/ml) lesions of the RSC. NMDA was injected by hand in volumes of 0.20 – 0.35 μl using a custom-made glass injection canula (100μm diameter) attached to a Hamilton Syringe by sterile plastic tubing. The stereotaxic coordinates and injection volumes were:

1. 0.35μL at −2.2 (AP), ±0.5 (ML), −3.0 (DV)
2. 0.35μL at −3.9 (AP), ±0.5 (ML), −3.0 (DV)
3. 0.20μL at −5.5 (AP), ±0.5 (ML), −3.5 (DV)
4. 0.35μL at −5.5 (AP), ±1.0 (ML), −2.8 (DV)
5. 0.35μL at −6.7 (AP), ±1.1 (ML), −2.8 (DV)
6. 0.30μL at −8.0 (AP), ±1.3 (ML), −2.8 (DV)

Coordinates were taken from Bregma (AP), the midline (ML), and the surface of the skull (DV), respectively. The injection cannula was left in place 1 min before and 5 min after each infusion. An additional ten rats received sham lesions of the RSC consisting of lowering the injection cannula into the brain but not injecting NMDA.

### Apparatus

Twelve rats were trained on a black PVC continuous T-maze (120 cm long stem x 100 cm wide x 68 cm above the floor) with metal reward cups embedded in the ends of the arms. Chocolate milk (0.2 ml, Nestle’s Nesquik) could be delivered to the reward cups via an elevated reservoir controlled by solenoid valves activated by foot-pedal switches. The maze was located in the center of a circular arena enclosed by black curtains with visual cues of various shapes, sizes, and colors. The room was illuminated by a ring of LED lights around the edge of the ceiling. A continuous background masking noise was played from a speaker located directly above the apparatus.

### Behavioral Training Procedures

Prior to training, rats were acclimated to the maze and chocolate milk rewards with daily periods of free exploration on the maze until rats consumed 20 rewards within the first 10 min of an acclimation session (mean = 4.5 acclimation days). After acclimating to the maze, rats were trained on a continuous spatial alternation task in which the rats were rewarded only if they approached the opposite (left or right) reward location from the previous trial. Both cups were baited on the first trial. Entries into the same arm as the previous trial were scored as an error and were not rewarded. Unlike some previous studies (43), rats were not shaped with trials where the incorrect choices were prevented by blocking access. Instead, rats were gently ushered back if they left the continuous alternation route. Rats were not allowed to correct their errors. Rats were given approximately 40 trials/day until they achieved a criterion of 90% correct on two consecutive days. After achieving this criterion, rats were given up to 10 additional training sessions to record neuronal activity during asymptotic performance (i.e. overtraining).

### Recordings

Neuronal spike data and video data were collected throughout learning (Digital Cheetah Data Acquisition System, Neuralynx, Inc. Bozeman, MT), filtered at 600Hz and 6kHz, digitized and stored to disc along with timestamps for offline sorting (SpikeSort3D, Nueralynx, Inc.). The rat’s position and head direction were monitored by digitized video of an LED array attached to the rat’s head. The time of reward receipt was measured with a grounding circuit that detected oral contact with the chocolate milk reward.

### Histology

After completion of the experiment, rats were transcardially perfused with 4% paraformaldehyde in phosphate buffered saline. Brains were removed and stored for at least 24hrs in 4% paraformaldehyde before being transferred to a 30% sucrose solution for storage until slicing. Coronal sections (40 μm) were stained with 0.5% cresyl violet for visualization of tetrode tracks (for neurophysiology recording implants) or tissue damage (in the case of NMDA lesions). Tetrode positions were identified using depth records noted during tetrode lowering and tracks observed in the stained tissue (**Supplementary Fig. 1**). Boundaries of the RSC were determined in accordance with The Rat Brain in Stereotaxic Coordinates (64). Neuronal records from tetrodes located outside of the RSC were excluded from the data set. As in our previous work (30), our recordings targeted the granular b subregion of the RSC, although small numbers of neurons from the granular a subregion or the dysgranular RSC were also included. There were no conspicuous differences in the firing properties of neurons recorded in different subregions, different hemispheres, or at different AP coordinates. Tissue damage was quantified by laying a grid (250 μm to-scale grid spacing) over an enlarged image of the stained tissue and dividing the number of grid intersections located over damaged RSC areas by the number of intersections located over the entire RSC. No significant relationship was seen between damage to different RSC sub-regions, or between extra-RSC regions, and alternation behavior.

### Data analysis

#### Spatial coding

Bayesian decoding was used to predict the current position of the rat on the maze given the spiking activity of simultaneously recorded RSC populations and a uniform prior (41) (**Fig. 1c**). This analysis was restricted to recording sessions with at least 8 RSC neurons (34 sessions). This population size was chosen to balance the trade-off between including only the largest populations and maximizing the number of included sessions (all population sizes are shown in **Supplementary Fig. 2**). Decoding was performed iteratively using a trial-based procedure whereby spike counts during time bins (200ms taken every 50ms) from one trial were used as the test sample, while the bins from all other trials were used as the training sample. The training sample was used to calculate firing rate maps for every neuron over a 50 × 50 grid overlaying the maze (mean of 352 visited pixels with each pixel approximately 2.5 × 2.5 cm). Probability distributions of spike counts for each neuron and pixel were computed based on the mean spike counts and assuming a Poisson distribution. For each time bin in the test sample, the probability of the rat being in a pixel was calculated by multiplying, across neurons, the conditional probabilities of observing those spike counts if the rat occupied that pixel. The highest probability pixel was taken as the decoded position of the rat on the maze, and was considered an instance of correct decoding if it was within 4.5 cm of the rat’s actual position (i.e. within a circle with a diameter of approximately half the body length of a rat). Decoding accuracy was compared to a distribution of chance accuracies obtained by shuffling 10,000 times the spike counts of each neuron independently among the time bins for each recording session in that learning stage. The observed accuracy was considered significant if it was greater than 97.5% of the shuffle outcomes.

Correlation matrices were created to quantify the selectivity and reliability of RSC spatial firing throughout learning (**Fig. 1e, f**). A single lap around the maze, beginning after the stem on a go-right trial and ending in the start area after a go-left trial, was divided into 170 spatial bins. Standardized mean firing rate vectors were then calculated for each spatial bin independently for the first and second half of each session (firing rate vectors contained the trial-averaged firing rate of every cell). To maximize comparability between learning stages, which had varying numbers of recorded neurons and systematic differences in behavior (i.e. more variable behavior was seen during early learning stages), the firing rate vectors were assembled from the first 50 neurons recorded during that stage and the analysis was limited to trials where the rat made typical passes through the maze section. Typical passes were defined as path lengths through a maze section that were shorter than 50% of all observed path lengths (from all rats and learning stages) through that section. This criterion was chosen because it eliminated instances where the rat’s trajectory through space was interrupted by backtracking, digressions, or pauses (see **Supplementary Fig. 3**). Sessions with fewer than five typical passes for each trial type (left and right) and session half were excluded. Separate correlation matrices were then generated for each learning stage (early, middle, late, and overtraining). Each row of a correlation matrix corresponds to the correlation of the ensemble rate vector for one spatial bin during the first half of the session with the ensemble rate vectors for every spatial bin during the second half of the session. If spatial firing was perfectly reliable between the first and second halves of the session, then the highest correlation would always be between a spatial bin and itself (e.g., bin 1 in the first half of the session would be most correlated with bin 1 in the second half of the session). In this case, the diagonal from the upper left to the lower right would contain the highest r value in each row. We therefore quantified spatial coding error by computing *divergence* from the diagonal. Specifically, mean spatial coding error was computed by summing, over all rows, the distance between the observed maximum correlation and the diagonal, multiplying this value by the length of each bin (3cm) and then dividing by the number of bins (170). To be conservative, the maze was treated as circular for computing distance, and the shorter of the two distances (forward or backward) between the reconstructed and actual positions was always used. Higher values indicated poor spatial coding. The observed spatial coding error was compared to a chance distribution of spatial coding errors computed by shuffling the first-half second-half neuron pairings. To then determine the statistical significance of differences between training stages we compared the observed differences (in terms of mean squared error, MSE) to a distribution of differences obtained by shuffling the 200 neurons (50 per learning session) randomly between the four stages 10,000 times and each time recalculating the total spatial coding errors for each stage and the MSE between them. The observed MSE was considered significant if it was greater than 97.5% of shuffle-generated MSEs. This analysis included only a single stem traversal (from the right to left reward location) for simplicity because trial-type specific firing on left and right trials can affect the correlations. However, similar results were obtained when the stem was included twice (as two separate trajectories for left and right trials) or when the stem was excluded.

#### Trial-type specific firing on the stem

Individual neurons exhibiting trial-type specific firing as the rat traversed the stem were identified by comparing firing between left and right trials in each of four equal sized stem sectors (see **Fig. 2a, Supplementary Fig. 5b**) using a two-way, repeated-measures ANOVA as in (43). Analyses were restricted to correct trials with stem runs that did not involve pauses or deviations from smooth locomotion (i.e. typical stem runs). Typical stem runs were defined as passes through the stem of the maze that took less than 1.24s. This criterion was 2.5 standard deviations above the mean run time and eliminated trials with irregular behaviors (e.g., backtracking, digressions, or pauses), and excluded 13.77% of learning trials and 3.63% of overtraining trials, see **Supplementary Fig. 5c**). To avoid comparing learning stages with different numbers of correct trials, only the first 10 correct trials of each trial type were included from each session. The statistical significance of the observed proportion of neurons was determined by shuffling 10,000 times both the (1) firing rates on each trial between the four sectors and (2) whether a trial was considered go-left or go-right, while maintaining the original proportion of each type. The observed proportion was considered significant if it was greater than 97.5% of shuffle proportions.

To assess the trial-type specificity of ensemble firing on the stem, we combined neurons across rats and sessions to form ensemble firing rate vectors during stem traversals on left and right trials, and then computed a specificity measure that quantified how similar activity during each stem traversal was to other traversals of the same trial type (e.g., left vs. left) and of the opposite trial type (e.g., left vs. right). Only the first 13 correct left and the first 13 right trials were included for comparisons over learning (**Fig. 2c-e**). The first 17 trials of each type were included for the overtraining-only analyses due to greater numbers of correct trials (**Fig. 2f-h**). To do this comparison, we used an iterative procedure whereby we excluded one trial from the data set, calculated mean left and right firing rate vectors from the remaining trials, and then computed the standardized Euclidean distance between the excluded trial and the two means. Specificity was then computed as the difference between the two distances normalized by the total distance. Positive specificity values (i.e. activity was more similar to the same trial type) were considered accurate classifications. Classification accuracy for each training stage (**Fig. 2e**; or behavioral performance, **Fig. 2h**) was compared to a chance distribution calculated by shuffling trial type labels 10,000 times, and the observed classification accuracy was considered significant if it was greater than 97.5% of the control classification accuracies. To determine whether the observed classification accuracies differed between the four training stages, we shuffled individual neurons between stages 10,000 times, and then calculated the MSE of the four control classification accuracies after each shuffle. The observed MSE was considered statistically significant if it was greater than 97.5% of the shuffled MSEs.

#### Reward location representations

We assessed the specificity of ensemble firing at the reward locations (**Supplementary Fig. 4**) in the same manner as the above analyses of ensemble responses on the stem, but with reward location as the category variable. We compared the time window 1-3s after lick detection on left and right trials, as this was when the rats were most still.

#### Decoding to Future Reward Areas

Analyses of reward representations during stem traversals were similar to the above Bayesian analysis of spatial coding except that we sought to determine the degree to which the two reward locations were represented in the population activity rather than the rat’s actual current position on the stem (**Fig. 3a-c**). The analysis only included correct trials and the test sample was restricted to time bins as the rat traversed the stem. For each trial, we calculated the decoded probability (i.e. decoding) that the rat was in the reward areas. Reward areas included both the reward locations and the portion of the approach arms after the choice point (see boxes in **Fig. 3b**). Most reward area decoding was at or near the reward locations, but substantial decoding was also seen along the arms. To determine whether the reward areas were overrepresented relative to other non-stem areas, we normalized the amount of decoding to the reward areas by their relative size (proportion decoding divided by proportion of pixels) and compared the observed value to chance (i.e. a uniform distribution, proportion of decoded probability is equal to proportion of total pixels; dotted line in **Fig. 3d**). The statistical significance of each stage mean was calculated by comparing the observed distribution of session means to a value of 1.0 using a Bonferroni-corrected one-sample t-test.

Representations of the two reward areas (left and right) were then compared to each other to determine whether the rat preferentially represented the reward area that it was about to approach (**Fig. 3e**). The difference between the representations (decoded probabilities) of the correct and incorrect reward areas were computed and then standardized by their sum (correct minus incorrect divided by the total). Positive values indicate a greater representation of the correct reward area, while negative values indicate a greater representation of the incorrect reward area. The statistical significance of each stage mean was calculated by comparing the observed distribution of session means to zero as above.

## Supplementary Figure Legends

**Supplementary Figure 1.**
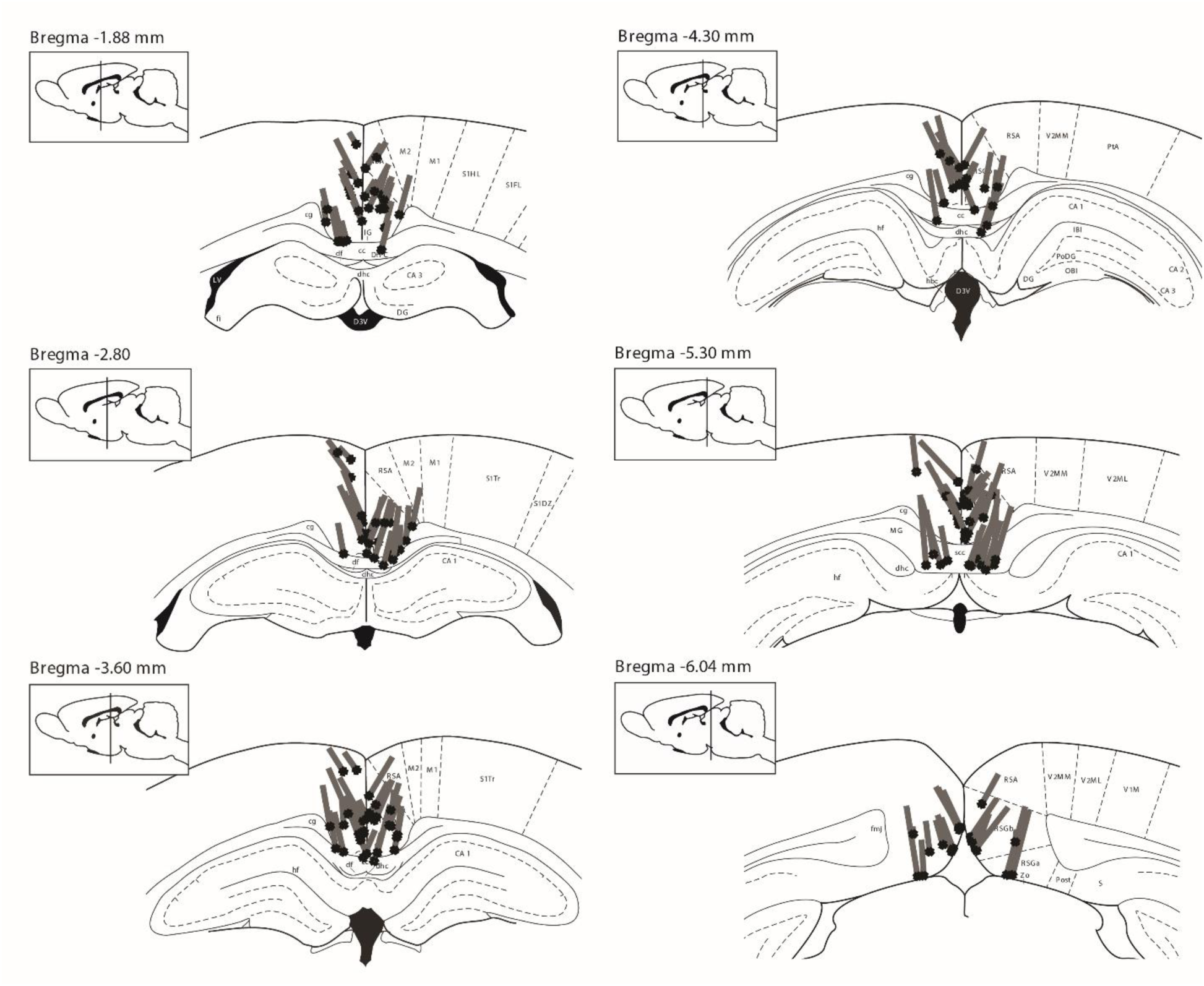
Estimated tetrode positions during recording (gray lines) and the corresponding end positions (black dots) are shown at 6 AP positions. End positions indicate the lowest point of the tetrode, and not necessarily the location of tetrode during the last recording session, which was often much earlier. Neurons were recorded primarily in the granular b subregion of the RSC (commonly referred to as Rgb, sometimes also referred to as Brodmann’s Area 29c), with smaller numbers of neurons recorded in the dysgranular and granular a subregions (Rdg and Rga, also referred to as Brodmann’s Area 30 and 29a&b, respectively). Any recordings from tetrodes located outside of the RSC were omitted from the analysis.

**Supplementary Figure 2.**
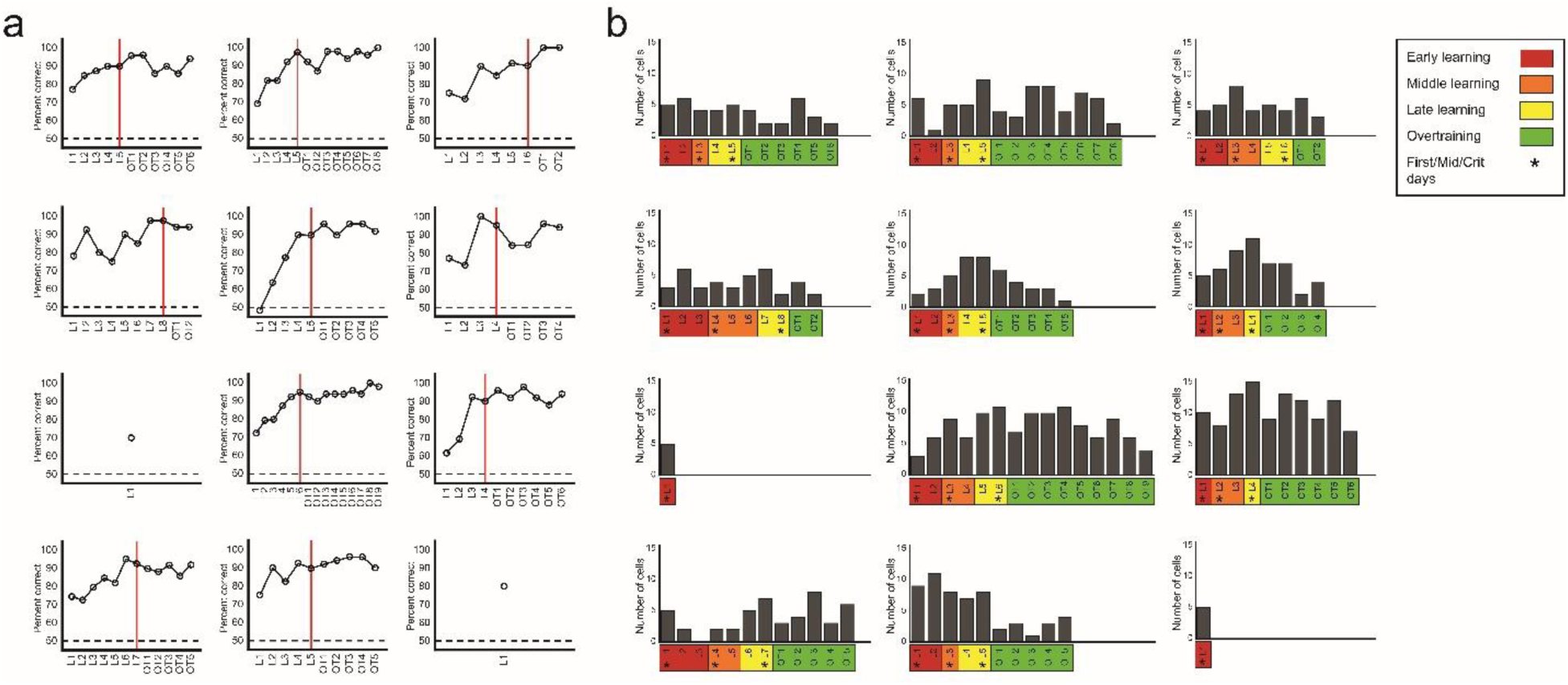
Learning curves and number of recorded cells for each rat included in the physiology analyses. (**a**) Rats received daily learning (L) sessions on the continuous spatial alternation task until they achieved a behavioral criterion of two consecutive days of at least 90% correct. The second day at or above 90% was considered the criterion day (vertical red line). Rats then received up to 9 overtraining sessions. The implants of subjects R1812 and R1861 were damaged before completion of training, and therefore only the first training day from each of these subjects is included in the data set. (**b**) Bar charts show the number of neurons and the specific training sessions that contributed to neural analyses for each subject. For most analyses, we used data from one specific session from each subject (First, Middle, Criterial sessions), and these are indicated by asterisks. However, we combined data from multiple sessions for some analyses that required additional neurons (e.g., analyses of simultaneously recorded populations). Combined sessions are indicated as Early (red), Middle (orange), or Late (yellow) training stages.

**Supplementary Figure 3.**
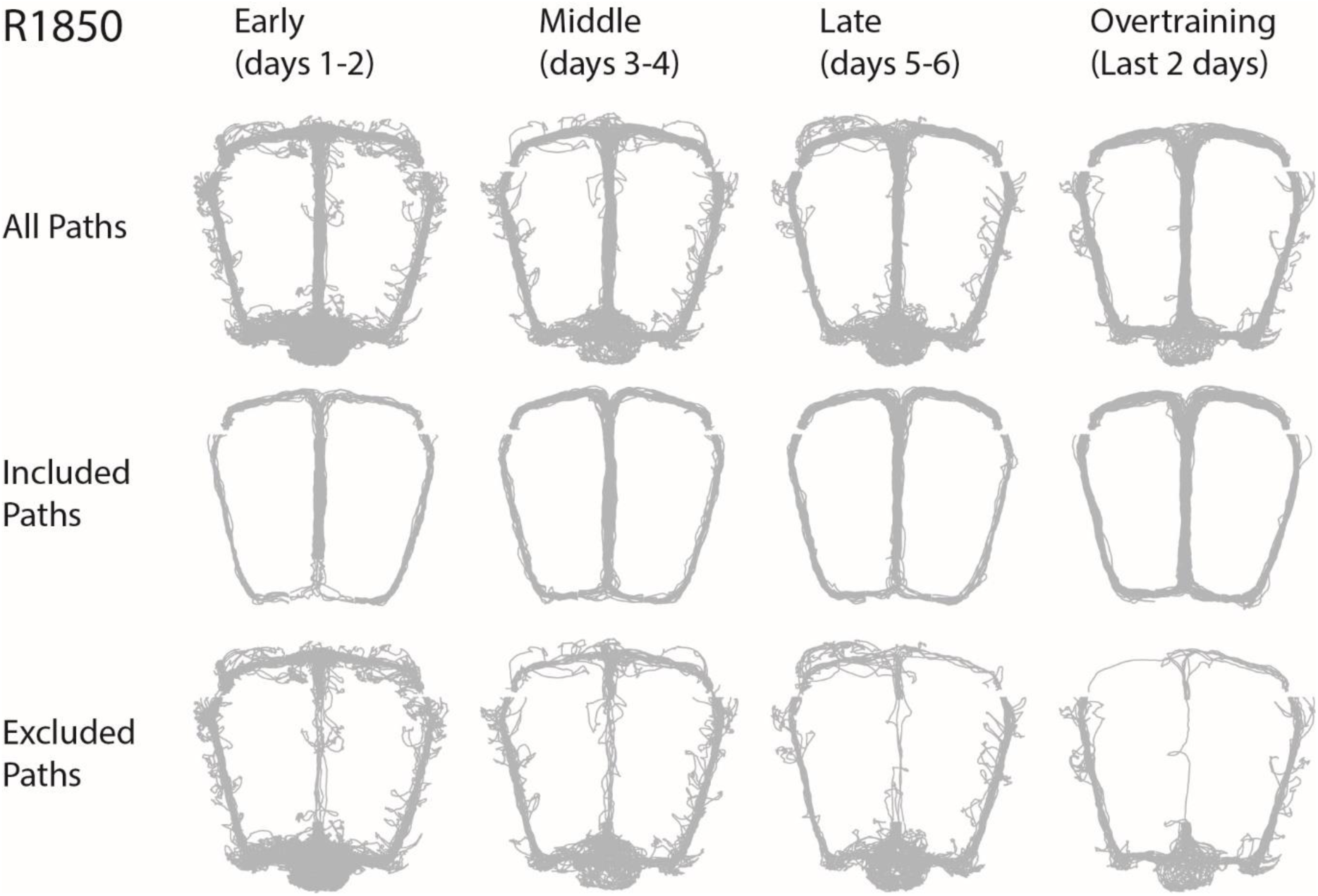
The path filtering process used to ensure similar behavior across learning stages. Grey lines in the shape of the maze show paths through space on every trial during Early, Middle, and Late learning stages, as well as Overtraining from one rat. Rats occasionally deviated from the most direct route along the maze, especially early in learning (top row). To ensure that we analyzed similar behavior at every learning stage, we separated the maze into 6 sections (start area, stem, left arm, right arm, left return, right return), combined paths across all learning stages, and excluded the longest 50% of paths through each section. This ensured that direct paths that were common to all learning stages (included paths, second row) were included, while excluding paths where the rat paused or backtracked (excluded paths, third row).

**Supplementary Figure 4.**
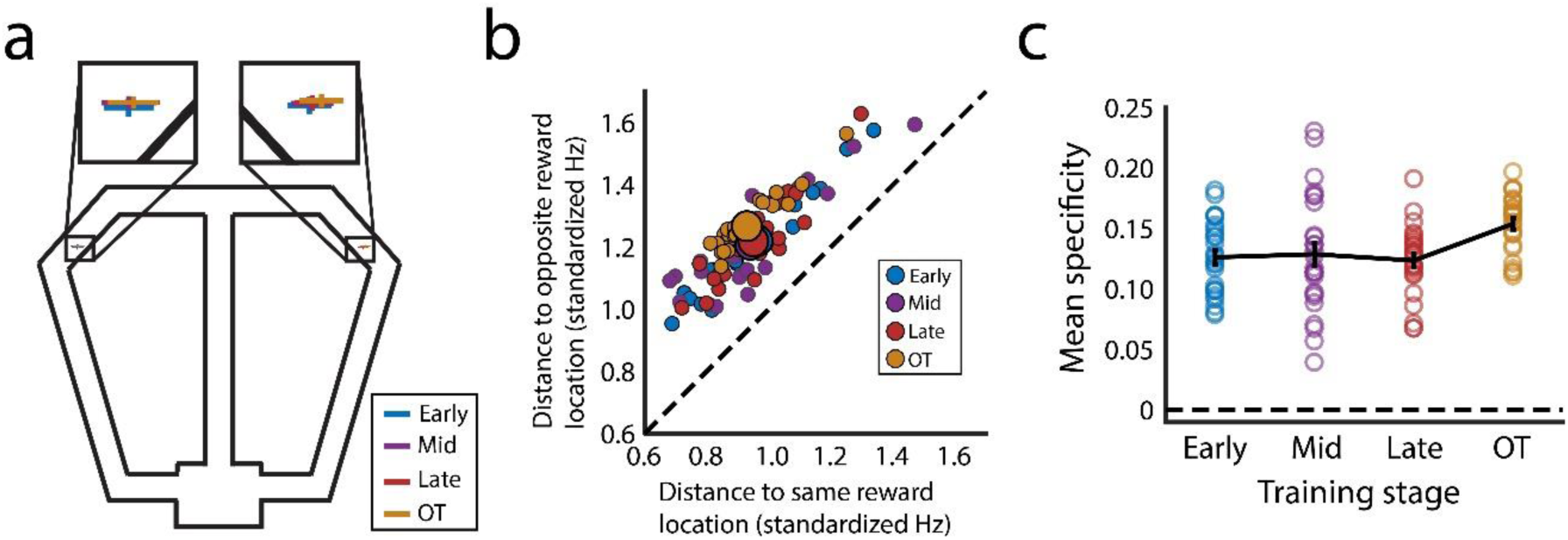
Behavior and ensemble activity at the left and right reward locations. (**a**) The position of the rats while they consumed the reward remained consistent throughout training. Mean positions +/- STD are shown for every learning stage at each reward location for the time window from 1 to 3 s after lick detection. (**b**) RSC ensemble activity (combined across subjects and sessions, see Methods) was distinct at the two reward locations throughout training. Each colored dot shows ensemble activity from one trial plotted in terms of its distance from the mean of the same and opposite reward location representations. Points along the dotted line are equidistant to both reward locations, while points farther from the dotted line indicate stronger ensemble preferences for one reward location over the other. Large dots with black outlines illustrate the mean for each learning stage. Note that ensemble activity moves further away from the unity line during overtraining (OT). (**c**) Ensemble activity during reward visits is plotted in terms of mean specificity, defined as the difference between the distances to the representations of the opposite and same reward location, standardized by the sum of both distances. Individual reward visits are plotted as open circles, with the mean for each training stage illustrated by the line plot +/- SEM.

**Supplementary Figure 5.**
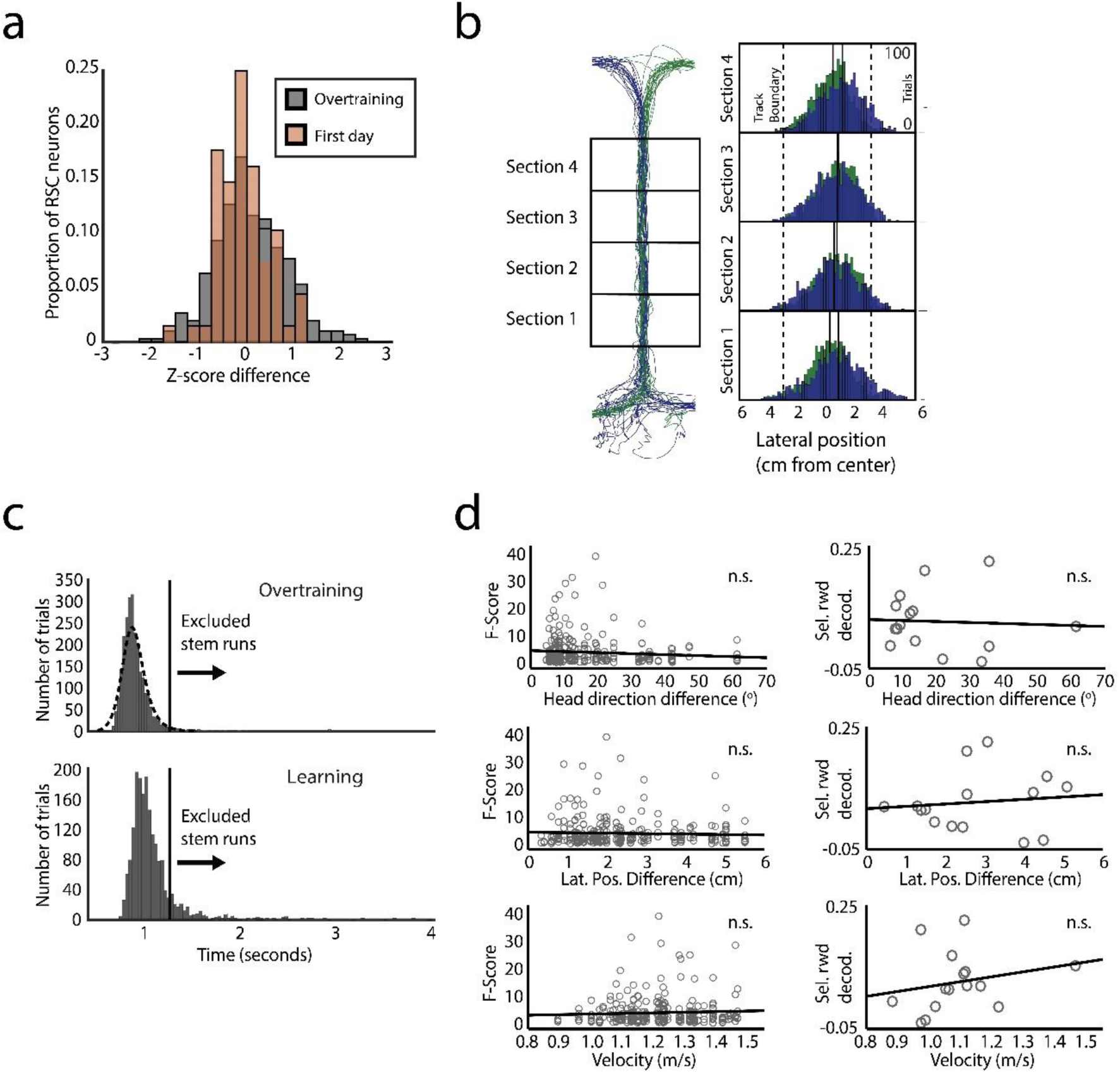
Behavior and trial-type specific firing on the stem of the maze. (**a**) Histogram illustrating the distribution of trial-type specific firing in individual RSC neurons. For each neuron, we calculated the difference between the mean firing rate of all right trials and all left trials, and then divided this by the pooled standard deviation (i.e. the z-score difference). Negative scores indicate higher firing on left trials, while positive scores indicate higher firing on right trials. The distribution on the first day (orange) is overlaid on the overtraining distribution (gray). (**b**) Stem positions on left and right trial types were highly overlapping in each of the four stem sectors used to define trial-type specificity in individual neurons (see Methods). All the included stem runs from one overtraining session are plotted (left), with left trials shown in blue and right trials shown in green. The lateral position on the stem of all overtraining trials in the dataset with left and right trials plotted for each sector (right). Bars of the histogram correspond to the number of trials where a rat occupied that lateral position on the stem, the solid black lines indicate trial-type mean position, and the dotted lines indicate the left and right boundaries of the stem. Because we determined the rat’s spatial position using lights on the head of the rat, the rat could be considered outside the stem boundaries if he tilted his head to the side. (**b**) Latency to traverse the stem was used to exclude trials with atypical behavior, with particularly high dwell times being indicative of pauses or backtracking. Histograms show the stem dwell times of every trial recorded during overtraining (top) and learning (bottom) sessions. To exclude atypical stem behavior, a log-logistic curve was fit to the distribution of all stem-run durations during overtraining (dotted line). The solid black lines indicate a cutoff at 2.5 standard deviations above the mean of the fitted curve. This cut off (1.24s) was then used to exclude stem runs from both overtraining and learning sessions. (**c**) Trial-type neural activity occurring on the stem of the maze was not correlated with head direction (top), lateral position (middle), or running speed (bottom) for either single-cells (left column) or simultaneously recorded populations (right column). Left column, circles show the relationship between neural activity and behavior for one cell. F-scores from the analysis for trial-type specificity in individual neurons (**Fig. 2b**; see Methods) were used as an indicator of trial-type specific neural activity. This measure was not correlated with behavioral differences on left and right trials. Right column, each circle shows reward-location specific decoding from one simultaneously recorded population from an overtraining session (**Fig. 3e**). This population activity was also not correlated with behavioral differences.

**Supplementary Figure 6.**
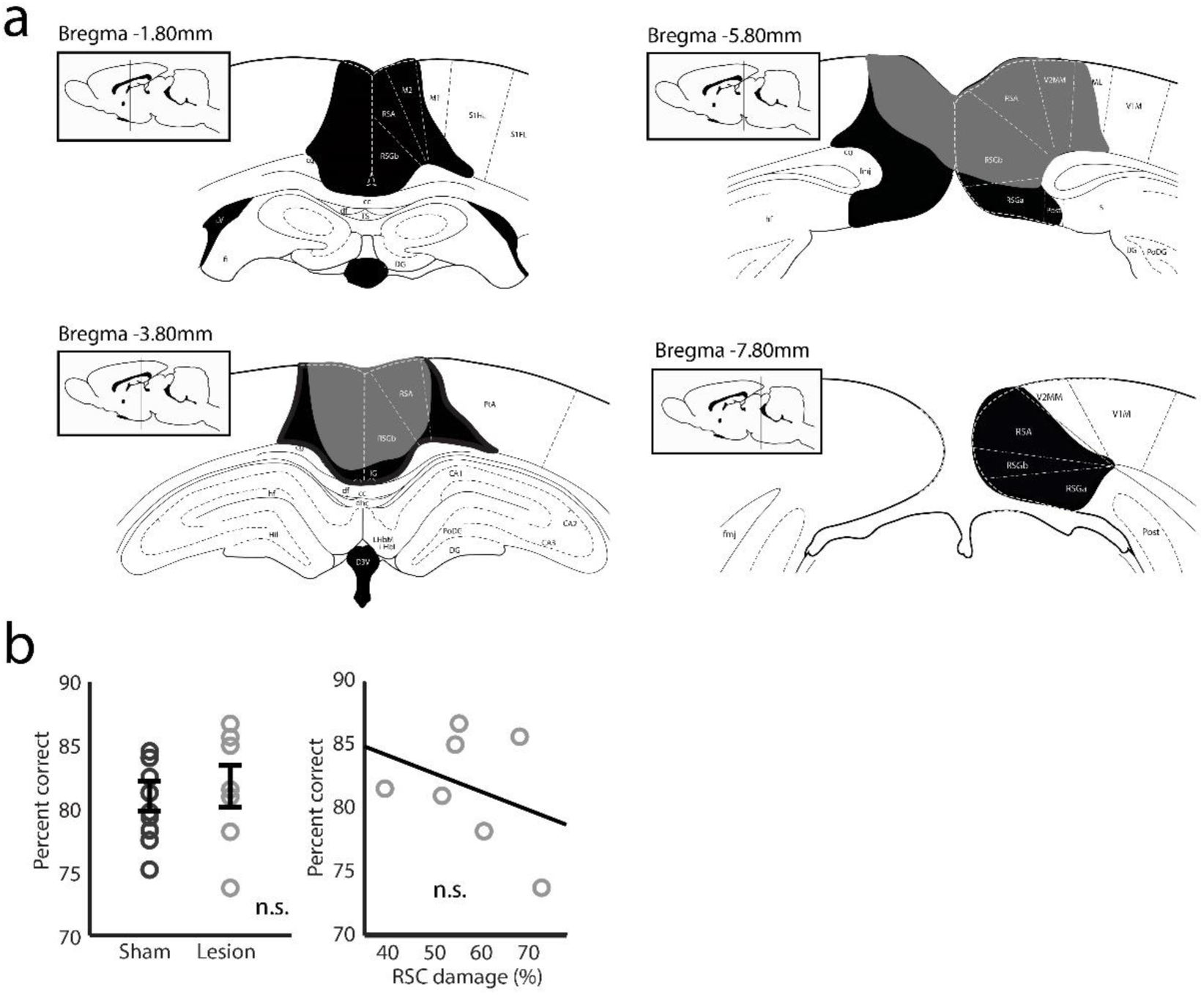
Lesions of the RSC. (**a**) Damaged area observed in the rats with the largest (black) and smallest (gray) lesions are shown at 4 AP positions. (**b**) Rats with lesions of the RSC showed similar alternation accuracy during learning (top, t_(15)_ = 0.33, p = 0.75). The correlation between alternation accuracy during learning and lesion size did not reach significance (bottom, r = −0.36, p = 0.43). Open circles correspond to individual rats’ mean percent correct over all learning days. Error bars show mean performance in each condition +/- SEM.

